# Fertilizers, BT technology, and insecticides contributed to 60%, 23%, and 17%, respectively to the increase in BT cotton yield: An analysis from 2000 to 2014 in India

**DOI:** 10.1101/2020.05.06.081794

**Authors:** C Parameswaran, B Cayalvizhi

## Abstract

Complementary technologies and agricultural practices capable of sustaining profitability to the farmers cultivating BtCotton in India require urgent attention. In India, approval of Btcotton, cultivation of fertilizer-intensive hybrids, higher dose of fertilizer application by farmers, usage of novel pesticides all happened simultaneously during the same period (2002-04) which makes very difficult to identify the individual effect in the yield gain of cotton. In this background, we attempted to understand the proportionate contribution of fertilizers, BT technology and novel group of pesticides in enhancing cotton yield in India. For the analysis, linear regression model and change in partial factor productivity (PFP) of cotton was considered in four different scenarios for yield estimation between 2000 and 2014, i.e. Scenario I: Cotton yield in the absence of technology and enhanced fertilizers usage, Scenario II: Cotton yield only due to enhanced fertilizer usage, Scenario III: Cotton yield with enhanced fertilizer and application of novel pesticides for the insect control, and Scenario IV: Cotton yield due to BT technology, enhanced use of fertilizer, and novel insecticides (actual yield of Cotton in India during Bt phase).Their comparison showed that the individual effect of fertilizers, BT technology and insecticides contributed to 60%, 23% and 17% of cotton yield, respectively in India. Further, 18% reduction in PFP was observed recently as compared to 2003-08. Besides, 125 Kg/ha of fertilizers was identified as optimum dose for sustaining high yield in cotton. Thus, present analysis identified the individual effect of different technologies contributing to the yield of cotton in India which can be used in decision making processes for crop improvement. Further, in our opinion, three strategies namely drip fertigation, intercrossing Bt and non-Bt hybrids for resistance management in bollworms, and IPM for sucking pests will primarily drive the research priorities and policy actions for the next 5 to 10 years in sustaining the economic benefits of the six million cotton farmers in India.

## Introduction

Cotton yield gain story in India is one of the inspiring technological achievements. In India, ∼ six million farmers cultivates cotton in an area of about 12 million hectares (2017-18) and approximately 50 million people are either directly or indirectly dependent on cotton for their employment and livelihood. Further, India is the leading producer of cotton in the world. However, yield of cotton in India requires further improvement in spite of more than two fold increase in the yield was achieved during the last two decades (Annual Report, AICRIP, 2017-18). Specifically, adoption of Bt cotton has coincided with the significant enhancement in the cotton yield in India (Whitefield, 2003).

India has succeeded in the major challenge which affected the cotton cultivation during 1990s due to the development of insecticide resistance in major pests of cotton (Armes et al, 1996; Kranthi et al, 2002).Especially, yield was stagnated or even reduced between the years 1995 and 2002 in India largely due to the resistance of cotton pests for pyrethroids and organophosphate groups of insecticides (Dhawan, 1998). In a turnaround, application of novel group of insecticides such as spinosad along with the adoption of integrated pest management has significantly increased the cotton yield in India during 2003-05 (Kranthi et al, 2000; Reddy et al, 2000, Kranthi, 2017).In the mean time, India has approved the cultivation of Bt cotton during 2002 (Jayaram, 2002). Further, impact studies of Bt cotton in India showed that yield increase was up to 19-24% in cotton, resulted in sustainable reduction in pesticide use, and also highlighted the contribution of fertilizers in yield enhancement(Kathage and Qaim, 2012, Krishna and Qaim, 2012, Gruere and Sun, 2012).In addition, Davis and Kranthi (2020) has recently reported that yield trend of cotton in India was highly correlated with the increase in the fertilizer application in India.

In India, more than two-fold increase in the cotton yield was observed between 2000 and 2015. Since, all the three practices namely adoption of Bthybrids, application of novel group of insecticides, and enhanced fertilizer usage by the farmers has happened simultaneously during the last 15 years in cotton, parsing out the individual effect of these technologies on yield gain is considered as relatively difficult (Personal communication from Prof. GD Stone). However, understanding the specific effect of technologies in yield enhancement on cotton is highly essential to devise the specific research and policy priorities for sustaining yield and profit of cotton farmers in India. In this regard, yield of cotton was affected by different factors in the last 30 years. For example, yield stagnation observed during 1996-2002 was majorly due to the wide-spread insecticide resistance developed in the cotton pests, whereas yield gain observed during 2003-05 was majorly attributed to the introduction of new class of insecticides. Further, combination of Bt technology and fertilizers has contributed to the significant yield increase in cotton after 2005. Thus, the aim of the present work was to comparatively analyze the impact of different technologies in four different time periods which was correlated with differential yield response in cotton.

Partial factor productivity (PFP) is the measure of yield per unit quantity of fertilizer applied (Wang et al, 2018). PFP is considered as one of the variable for understanding the long-term yield trends in crops (Fixen et al, 2015). For example, rate of change in the nitrogen-partial factor productivity in rice between 1980 and 2014 was attributed to optimum input management and mechanization in northern China (Zhang et al, 2017). Moreover, cotton yield in India was also characterized by varying PFP from 2000-2014 (Stone and Kranthi, 2020). Therefore, we hypothesized that the change in partial factor productivity of cotton could effectively predict the specific impact of different technologies such as Bt hybrids, fertilizer and insecticides influencing the yield gain of cotton in India.

## Methodology

### Yield trend in cotton

The relationship of fertilizers and Bt Cotton area with that of the cotton yield was studied using linear regression model(Supplementary Table 1). The analysis showed that both the fertilizers and Bt cotton area were highly correlated to the cotton yield with high R^2^ value(Supplementary Figure 1,2). The impact of fertilizers on the yield of cotton was previously reported by Kranthi and Davis, (2020). However, reasons behind the rapid adoption of Btcotton in India requires further investigation. Thus, the present work initially hypothesized that Bt technology has also contributed to the yield gain in cotton additionally to fertilizers. But, determining the individual impact of Bt technology, fertilizers and insecticides to the cotton yield will essentially requirea time-scale analysis of yield trend in cotton, i.e. before and after the introduction of Bt cotton or before and after the introduction of novel insecticides for the control of bollwormin India. Therefore, in order to determine the individual effects of technologies (Bt cotton and novel group of pesticides) and management (fertilizers) practices on the cotton yield trends, four different scenarios were considered (Table 1).

**Table 1.** Four different scenarios for understanding the change in yield trends in cotton

| Scenarios | Time Periods-Initial Phase | Time Periods-Later phase | Conditions during initial phase | Conditions during later phase | Reasons |
| --- | --- | --- | --- | --- | --- |
| **Scenario I**<br>(Yield estimation in absence of any technology) | 1991-1995 | 1996-2003 | Increasing yield trends | Decreasing yield trends | Decreasing yield trend was due to wide-spread insecticide resistance observed in bollworms |
| **Scenario II**<br>(Yield estimation only due to the fertilizers) | 2000-2003 | 2004-2014 | Stagnant yield | Increasing yield trends | Stagnant yield was due to wide-spread insecticide resistance observed in bollworms and increasing trends was due to fertilizers, adoption of Bt and novel insecticides in the control of sucking pests |
| **Scenario III**<br>(Yield estimation due to the fertilizers and novel insecticides) | 2003-2005 | 2006-2014 | Yield gain | Increasing yield trends | Yield gain due to effective control of bollworm by novel group of insecticides, other practices and increasing yield trends was due to fertilizers, adoption of Bt and novel insecticides in the control of sucking pests |
| **Scenario IV**<br>(Yield estimation due to the Bt, fertilizers and insecticides) | 2005-2008 | 2009-2014 | Increasing yield trends | Increasing yield trends | Increasing yield trends was due to fertilizers, adoption of Bt and novel insecticides in the control of sucking pests |

The estimation of cotton yield in different scenarios are given below,

#### a) Estimation of yield in Scenario I

Cotton yield in India between 1996 and 2003 (Supplementary Table 2) was almost stagnant due to the development of insecticide resistance in bollworms especially for synthetic pyrethroids. Thus, extrapolation of yield trendsobserved during 1990-2003 will effectively predict theyield of cotton from 2003 onwards in absence of an effective technology to control the pests. Linear regression model was used to understand the yield trends between 1990 and 2003 (Supplementary Fig. 3). The linear regression equation for yield is given below,

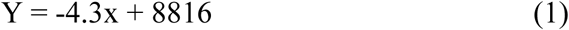

where, Y is yield of cotton, and x is year (1990-2003). Besides, R^2^ value for the linear regression model was found to be 0.46. Further, thisregression equation (Eq.1) was used to extrapolate the yield of cotton between 2004 and 2014 (Supplementary Table 3).

#### b) Estimation of Yield in Scenario II

Partial factor productivity, a measure of the yield per unit quantity of fertilizers applied (Wang et al, 2018) was used to estimate yield in Scenario II using the formula given below,

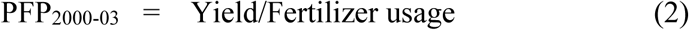

where, PFP_2000-03_ is partial factor productivity from 2000 to 2003. The determination of PFP for the pre-Bt period (2000-03) provides us a reliable prediction of yield gain which could have been achieved specifically due to application of fertilizers in the absence of Bt technology and insecticides. Besides, Scenario II presumes no effective insecticides were available for the control of pests in cotton. Moreover, PFP of cotton was almost stagnant from 1996 to 2003 in India (Suresh et al, 2014) indicating the yield stagnation during this period. The analysis showed that PFP_2000-03_was 2 Kg/ha during the pre-Bt period (2000-03). Further, PFP_2000-03_ was used to estimate the yield in Scenario II from 2004 to 2014 using the formula given below,

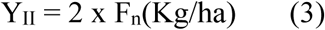

where, Y_II_ is the yield of cotton in Scenario II, F_n_ is the actual fertilizer used per hectare for the years between 2004 and 2014. Additionally, high incidence of sucking pests observed after 2008 due to heavy application of fertilizers has resulted in yield reduction in cotton. Therefore, yield loss due to sucking pest was also considered for calculating the yield in Scenario II. For determining the yield loss due to sucking pest, reduction in PFP from 2009 to 14 was initially estimated using the formula given below,

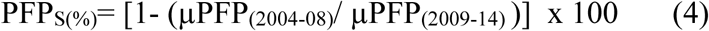

where, PFP_S(%)_ is the percent reduction in PFP due to high incidence of sucking pests, µPFP_(2009-14)_ is mean PFP from 2009 to 2014 and µPFP_(2004-08)_ is mean PFP from 2004 to 2008 during which the incidence of sucking pest was comparatively less. This estimate of reduction in the PFP_S_ was used to determine the yield in Scenario II from 2009 to 2014 using the equation given below,

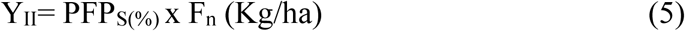

The estimated yield in Scenario II is given in Supplementary Table 4.

#### c) Estimation of Yield in Scenario III

The usage of pesticides having the novel mode of action for control of bollworm has dramatically increased during 2003-05 (Supplementary Table 5). Therefore, PFP for these two years also showed a significant increase over the PFP estimated from 2000 to 2003. Thus, increase in PFP during 2003-05 might have been attributed to the effective control of cotton pests using novel groupsof insecticides. Besides, the difference in the fertilizer use was only 10 Kg/ha during 2003-05 as compared to previous years (2000-03). Thus, increase in the PFP observed during 2003-05 could be taken for estimating the cotton yield in Scenario III specifically due to the application of novel class of insecticides. The percent increase in the PFP due to the application of novel insecticides was calculated using the equation given below,

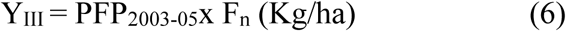

where, Y_III_ is the yield in Scenario III, PFP_2003-05_ is the partial factor productivity for the years during 2003-05 and Fis the actual fertilizers applied during the corresponding years. Further, yield loss due to sucking pest incidence was determined using the formula,

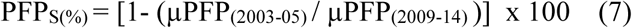

where, PFP_S(%)_ is the percent reduction in PFP due to high incidence of sucking pests, µPFP_(2009-14)_ is mean PFP during 2009-14 and µPFP_(2003-05)_ is mean PFP from 2003 to 2005 during which the application of insecticides significantly increased the yield of cotton (Supplementary Table 6).Finally, estimated reduction in the PFP_S_(%) due to sucking pest was used to determine the cotton yield in Scenario III after 2008 onwards using the below mentioned equation,

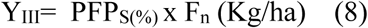

where, Y_III_ is the yield in Scenario III, PFP_s(%)_ is the percent reduction in partial factor productivity from 2009 to 2014 and Fis the actual fertilizers applied during the period. The estimated yield in Scenario III is given in Supplementary Table 5

#### d) Estimation of Yield in Scenario IV

The actual yield obtained due to adoption of Bt cotton was considered as the yield in Scenario IV.

### Percent contribution to yield by fertilizers, Bt technology and insecticides

The estimated yield of cotton in different scenarios was compared to understand the individual contribution of fertilizers, Bt technology and novel class of pesticides. The percent contribution of fertilizers contributing to cotton yield was estimated by

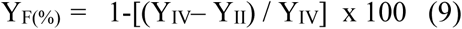

where, Y_F(%)_ is the percent yield contributed by the fertilizers, Y_IV_ is the actual yield of cotton due to Bt technology, fertilizers and insecticides, Y_II_ is the estimated yield in cotton specifically due to fertilizers. Similarly, yield contributed by the combination of Bt technology and insecticides was determined using the formula given below,

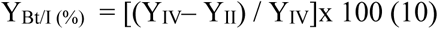

where,Y_Bt/I(%)_is the percent contribution to cotton yield by combination of Bt technology and insecticides,Y_IV_, and Y_II_ are estimated yield in different scenarios. Further, percent contribution of cotton yield specifically by adoption of Bt technology and insecticides was determined indirectly using the amount of insecticides used for control of sucking pests during 2004 and 2005. The pesticide quantity (%) used for the control of sucking pests between 2004 and 2005 was determined by,

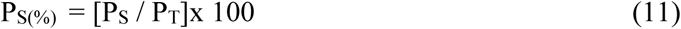

where, P_S(%)_ is the percent pesticide usage for the control of sucking pests, P_S_, and P_T_ is the quantity of pesticides used for the control of sucking pest and total pesticide used for the control of all the pest in cotton, respectively. Using the above equation, percent yield contribution of insecticide in the cotton yield was determined by,

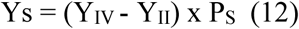

where, Y_IV_ is the yield in Scenario IV and Y_II_ is the yield in Scenario II, P_S_ is the percent pesticide usage for the control of the sucking pests. Finally, percent contribution by Bt technology was determined by

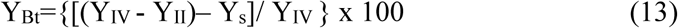

where, Y_Bt_is the yield contribution due to Bt technology.

### Statistical analysis

Analysis of variance (ANOVA) for the estimated yield in different scenarios was analyzed using the data analysis option in the MS Excel. Similarly, Z-test for the mean yield in different scenarios was analysed for statistical significance. For the analysis, null hypothesis considered was: no significant difference in the yield of cotton in four different scenarios. The test statistic and P value@ 5% and 1% was analysed for evaluating the null hypothesis and statistical significance, respectively.

## Results

The linear regression equation estimated for understanding the relationship between cotton yield and fertilizers was, Y = 0.356x + 15.92, where ‘Y’ is the yield and ‘x’ is the year. Further, R^2^ value was found to be very high (0.85). Similarly, linear regression equation of cotton yield and Bt cotton area also showed a high R^2^ value (0.87) (Supplementary Figure 1, 2). Additionally, multiple linear regressionsmodel for cotton yield with two variables namely fertilizers and Bt cotton areahasshowed relatively higher R^2^ value (0.92)and the obtained linear equation was, Y = 130.7 + 0.97a + 1.73b, where ‘Y’ is the yield of cotton, ‘a’ is fertilizer in Kg and ‘b’ is percent Bt cotton area in India. The initial yield trend analysis has identified that both the fertilizers as well as Bt cotton area highly influenced the cotton yield in India between 2000 and 2014.

Yield data of cotton during 1991-2003 was used to estimate the yield in scenario I. The analysis showed that the yield of cotton has actually reduced from 260 kg/ha in 1996 to ∼200 kg/ha for the years between 2000 and 2003,which indicates 23% reduction in the yield. Further, this decline in the yield was responsible for decreasing yield trend in cotton with low R^2^ value (0.54) between 1991 and 2003 (Supplementary Figure 3). This linear equation (y = - 4.3x + 8815, where ‘x’ is year) which showed reducing yield trend was used to extrapolate the yield of cotton in Scenario I for the years between 2004 and 2014. The analysis showed that the cotton yield has reduced up to 29% in 2014 (149 kg/ha) relative to the actual yield in the year 2000 (209 kg/ha). Since the Scenario I consideredthe widespread resistance of bollworm to insecticides, reduction in the estimated yield for the years 2004 to 2014 has followed the decline in the yield trend observed during 1996-2003 owing to the increase in damage caused by bollworm and other pests (Supplementary Table 3).

Partial factor productivity (PFP) was estimated for the years between 2000 and 2014 (Table 2). The analysis showed that the PFP of cotton has increased on an average of about 31%, 20% during 2004-08 and 2009-14, respectively as compared to the base year 2000. Thus, our analysis showed initial increase in PFP up to 2008 and thereafter mostly reducing trend was observed. Specifically, high incidence of sucking pest might be one of the major reasonsbehind the reduction in the PFP of cotton. Therefore, mean PFP of cotton during the years 2000-03 was used as reference for estimating the yield of cotton in the Scenario II. The PFP in cotton during 2000-03 was found to be 2 Kg/ha. Further, yield was estimated for the scenario II using the amount of fertilizers used for the cultivation. Additionally, high incidence of sucking pest has reduced the PFP by 17.6% from 2008-2014. Thus, the estimated yield in Scenario II was calculated after reducing the yield loss due to sucking pests. The analysis showed that the yield in scenario II was increased by 40% in 2014 due to the application of higher dose of fertilizers (Supplementary Table 4).

**Table 2.** Partial Factor Productivityfor cotton in India

| Year | Observed<br>Yield<br>(Kg) | Fertilizer<br>Usage<br>(Kg) | PFP<br>(Yield/Unit<br>qt) |
| --- | --- | --- | --- |
| 2000 | 200 | 100 | 2.0 |
| 2001 | 190 | 95 | 2.0 |
| 2002 | 190 | 95 | 2.0 |
| 2003 | 190 | 95 | 2.0 |
| 2004 | 290 | 100 | 2.9 |
| 2005 | 300 | 110 | 2.7 |
| 2006 | 350 | 120 | 2.9 |
| 2007 | 380 | 125 | 3.0 |
| 2008 | 410 | 130 | 3.2 |
| 2009 | 400 | 160 | 2.5 |
| 2010 | 400 | 160 | 2.5 |
| 2011 | 495 | 200 | 2.5 |
| 2012 | 480 | 200 | 2.4 |
| 2013 | 500 | 215 | 2.3 |
| 2014 | 460 | 200 | 2.3 |

Our analysis showed that between 2004 and 2005, PFP has increased by 29% relative to the base year 2000 (Table 1). This increase in the PFP was attributed to the application of novel group of insecticides for the control of boll worm. Therefore, percent increase in PFPobserved during 2004-05 was used to estimate the yield of cotton in Scenario III. Besides, as compared to 2005-08, sucking pest incidence has reduced the PFP by 21% during 2009-2014. Thus, 29% increase in the PFP due to insecticides and 21% reduction observed due to sucking pests were used in estimating the yield in Scenario III. The analysis showed that the actual yield in 2014 has increased up to ∼450 Kg/ha as compared to 200 Kg/ha observed during the base year 2000(Supplementary Table 5). Therefore, the estimated yield has increased up to 2.24 fold in scenario III during 2014. Similarly, actual yield has also increased up to 2.3 fold in scenario IV during the year 2014 as compared to 2000.

The comparison of yield in different scenarios showed that the combination of scenarios III and IV has increased the yield of cotton by 129 Kg/ha/year as compared to Scenario II (Fig.1). Additionally, increase in the yield of ∼99 Kg/ha/year was contributed specifically in Scenario IV as compared to Scenario III. Moreover, Scenario II has contributed to a maximum yield enhancement of 276 Kg/ha/year in comparison with reference year 2000. Thus, the percent increase in yield due to scenario III and IV is 40.4% out of which around 23.1% was contributed specifically in Scenario IV (Fig.2,3). Further, Scenario II has contributed to ∼60% yield enhancement in cotton. Besides, statistical test performed through analysis of variance (ANOVA) showed that there was a significant difference in the estimated yield between the different scenarios (P value: 1.26 × 10^−6^). However, estimated yield in Scenario III and IVdidn’t show the statistical significance (P value: 0.32). Additionally, significant difference was observed for the mean yield between the Scenario II and III (P value: 0.0059), and Scenario II and IV (P value: 0.0022).

**Figure 1.**
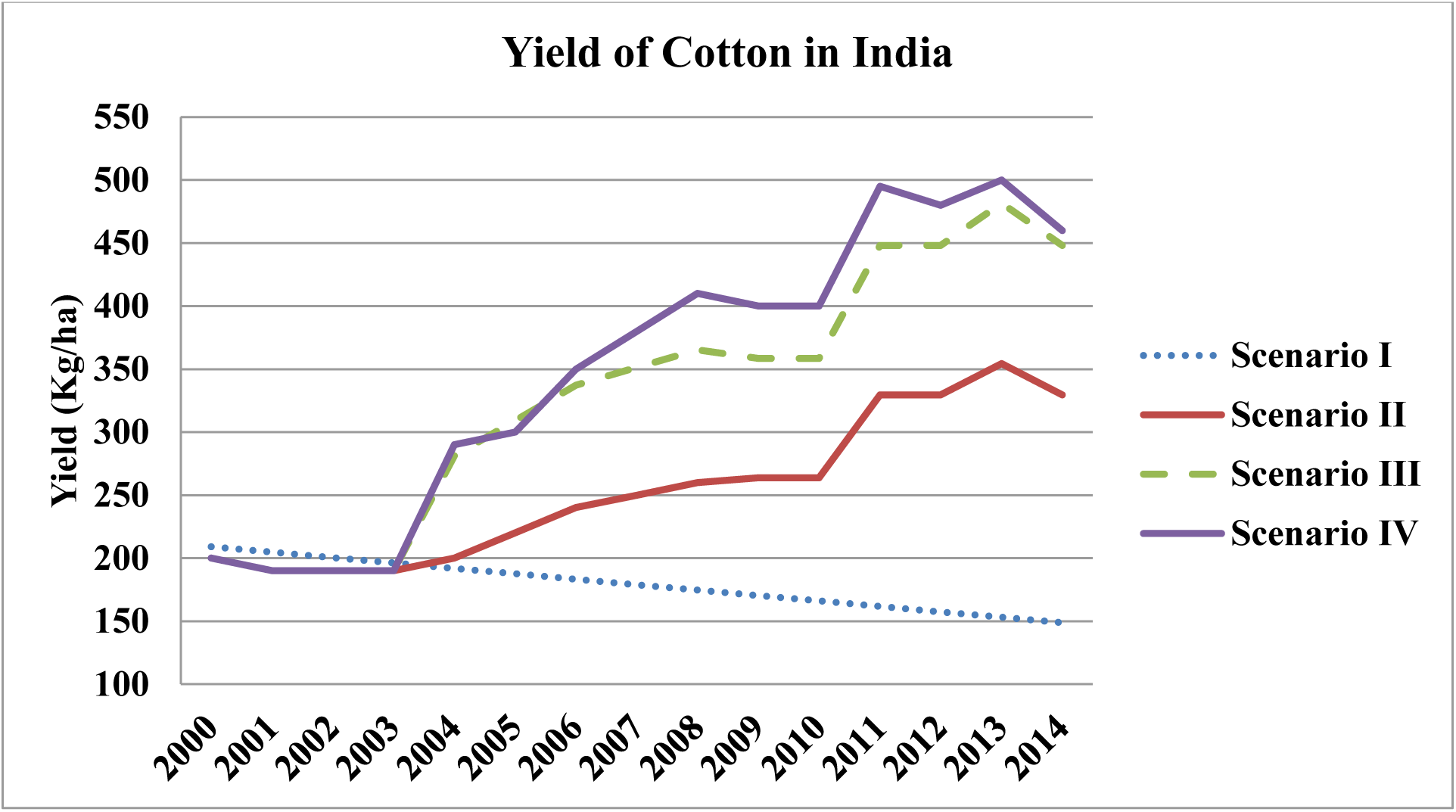
Yieldincrease in cotton estimated in four different scenarios

**Figure 2.**
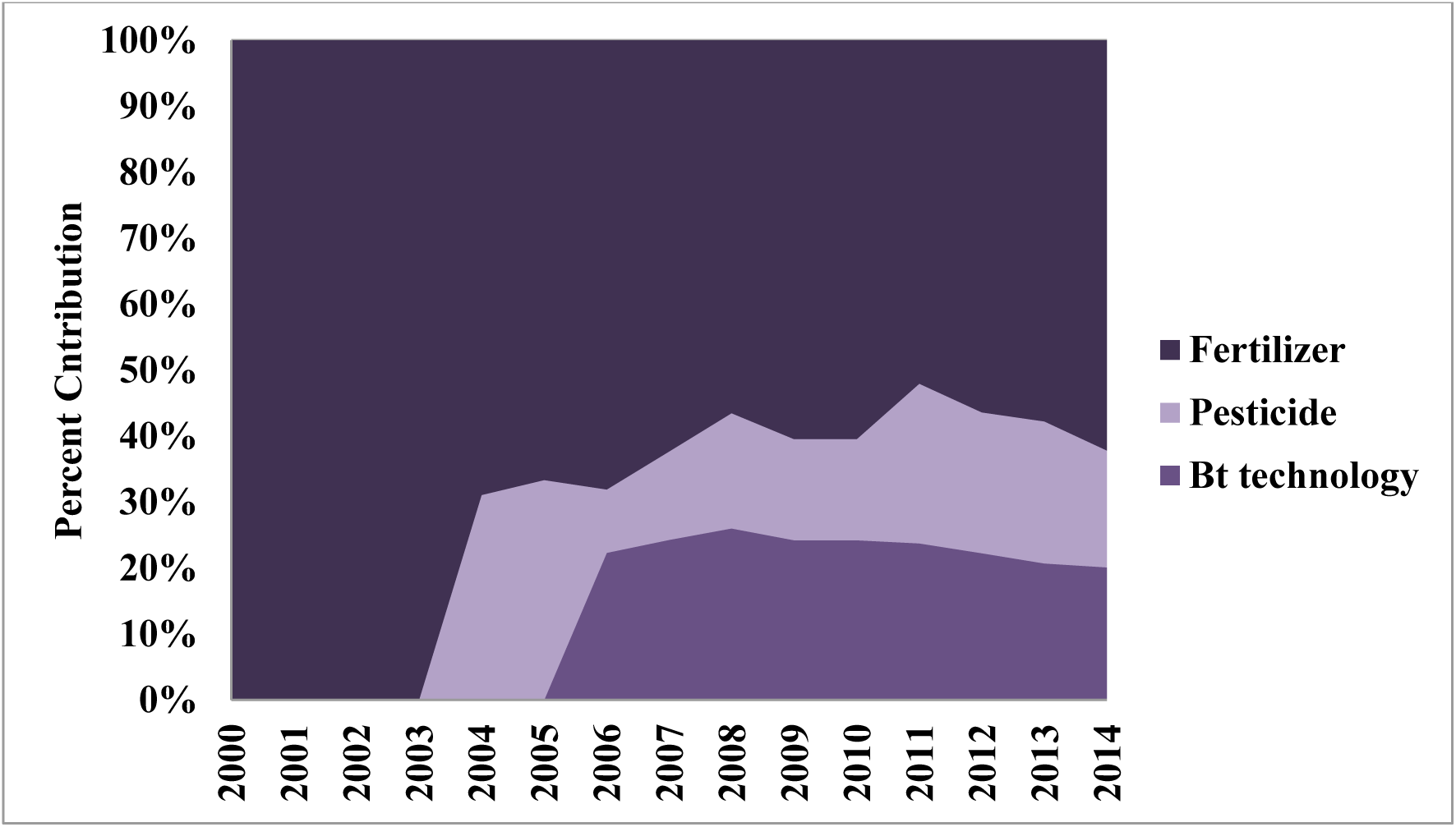
Effect of fertilizers, Bt technology and insecticides on the cotton yield in India

**Figure 3.**
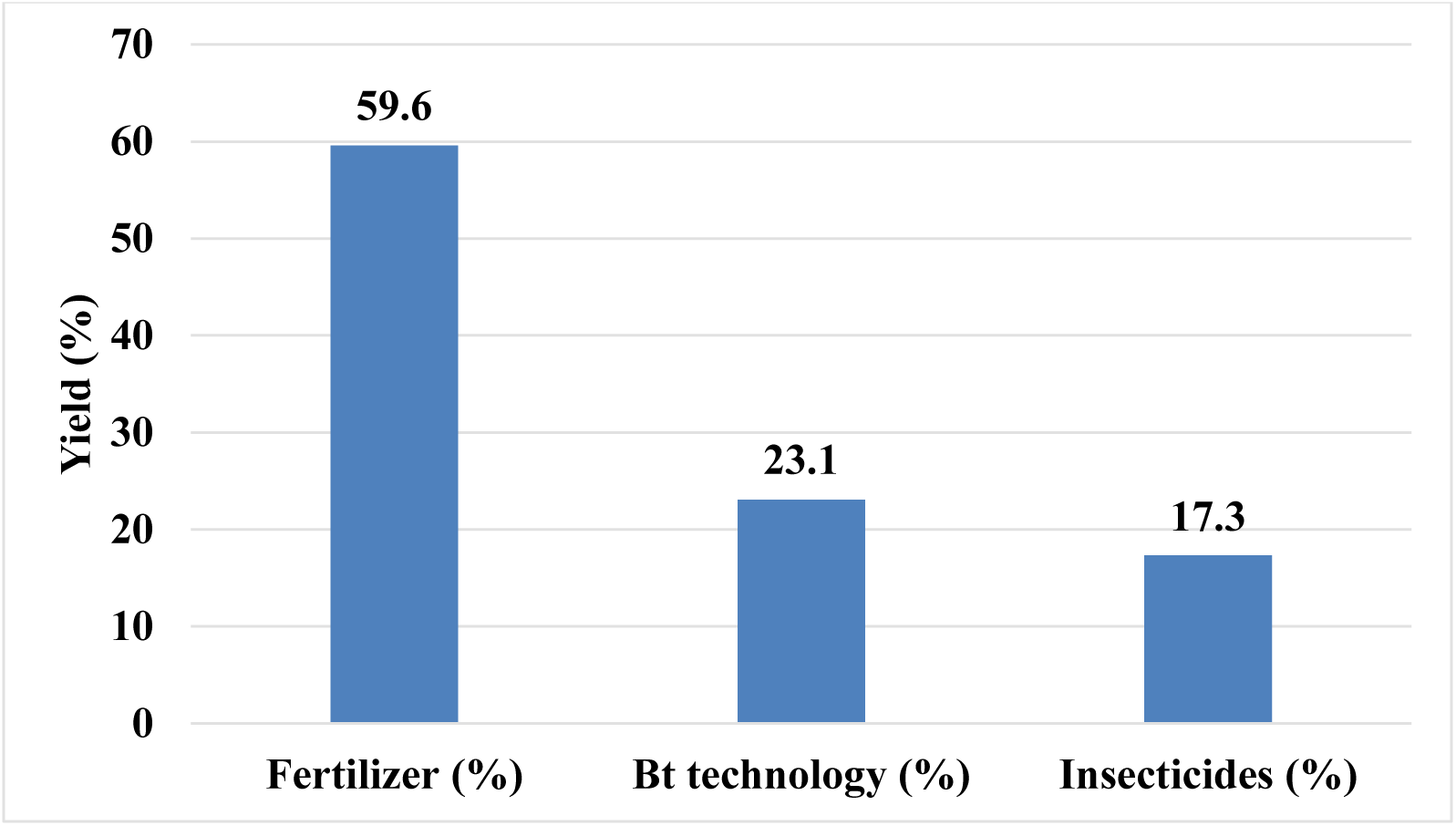
Contribution of fertilizers, Bt technology and insecticides on the cotton yield in India

## Discussion

The first major findings of our study is fertilizers contributed to an increase of about 59% in the yield of cotton as compared to 2000-01.There are several reports which showed increased fertilizer application @ 125% to the recommended dose of fertilizers (RDF) as well as closer spacing has increased the yield of cotton (Reddy and Gopinath, 2008; Bhalerao and Gaikwad, 2010; Bhalerao et al, 2012) in BT hybrids. Recently, Davis and Kranthi (2020) highlighted that yield trend in cotton was highly correlated with the increased fertilizer application. However, none of the previous studies could determine the proportionate contribution of fertilizers in increasing the cotton yield in India during the BT phase. Besides, BT cotton hybrids showed better nitrogen use efficiency and also responded well to the increased N application as compared to non-BT hybrids (Gangaiah et al, 2013). Therefore, increase in the yield of cotton in India during the BT phase could be due to the cultivation of nitrogen responsive Bt hybrids. However, response of cotton yield to Napplicationwas found to have quadratic relationship and showed maximum peak @ 180kg/ha (Gangaiah et al, 2013). Besides, higher dose of fertilizers has been reported to increase the population of sucking pests in cotton (Kalaichelvi, 2008).Thus, the observed reduction in PFP of cotton from 2009 to 2014 in India might be due to the higher dose of fertilizer application, i.e. 189 Kg/ha fertilizer was applied between 2009 and 2014 as compared to 125 kg/ha from 2005-08. Even though, yield of cotton in India is relatively less (500 kg/ha) as compared to China (1330 Kg/ha) and USA (900 kg/ha), continuous application of higher dose of fertilizers may not result in sustainable increase in the yield of cotton in India. In this regard, an optimum dose of 125 Kg/ha of fertilizersalong with drip fertigation (Wang et al, 2018) and biofertigation (Jayakumar et al, 2015) could become an effective practice in cotton cultivation for achieving better water, nutrient use efficiency, and sustainable yield improvement.

The second major findings of our study is BT technology has contributed to ∼23% yield gain in cotton over the reference year 2000. Even though BT trait cannot directly increase the yield of cotton, BT protein reduce the damages caused by bollworm and indirectly enhances the yield of the crop. Moreover, previous study from Kathage and Qaim (2012) also reported a yield increase of 24% during 2002-08 was due to Bt cotton in India. Besides, a meta-analysis of GM technology in multiple crops also showed that insect resistance in crops has increased the yield of different crops by 24.8% (Klumper and Qaim, 2014). Further, the proportion of yield gain achieved due to the cultivation of Bt cotton variedfordifferent countries and India, Argentina, Pakistan, Myammar and South Africarecorded a yield gain of >20% (Brookes and Barfoot, 2017).Additionally, economic benefits to the farmers obtained due to Bt cotton cultivation continued for 15 years since its initial adoption in China (Qiao, 2015). Thus, Bt cotton can be considered as the major technological achievement which has been benefiting more than 6 million farmers in India. However, high incidence of pink boll worm was recently reported in Bt cotton field in India (Naik et al, 2020; Mahesh and Mohan, 2020). In this regard, approaches such as releasing sterile moths, and planting of cotton hybrids developed through the cross between Bt and non-Bt cotton can be employed for the management of *Cry* genes resistant bollworms (Wan et al, 2017; Tabashnik and Carriere, 2019). In addition, RNAi technology for disrupting hormone metabolism in insects and combination with Bt technology was reported to be effective in controlling resistant bollworms (Ni et al, 2017). Thus, varied research priorities are essential for the effective management of bollworm resistance developed in the Btcottonin India.

The third major finding was insecticides has contributed to 17% yield gain in cotton. Even though, insecticide use for the control of boll worm has significantly reduced in India, sucking pest dynamics showed a significant change only after wide-spread adoption of Bt cotton (Bhute et al, 2012). Moreover, application of insecticides for the control of sucking pests has increased to more than three fold in India during 2011-12 as compared to 2002-03. Several previous reports showed that the application of insecticides against sucking pest has resulted in enhanced yield in cotton (Agale et al, 2010; Kumar et al, 2015; Nemade et al, 2017;Meena et al, 2018). In this scenario, increased spraying of insecticides by the farmers due to altered sucking pest dynamics have also contributed indirectly to the yield increase in cotton through minimizing damage caused by sucking pests. Moreover, a report by Oerke and Dehne (2004) has showed that the efficacy of pesticides was found to be high in the cash cropshighlighting the prime reason for increased insecticide application by the farmers. Besides, previous studies showed the influence of weather parameters were highly related to the increased incidence of sucking pests in cotton (Babu and Meghwal, 2014; Roomi, 2016; Chauhan et al, 2017).Thus, there is a growing concern that sustainable benefits of Bt cotton achieved during the last 15 years in India may be affected in future for the reasons of bollworm resistance to Bt, climate change, and high incidence of sucking pests. In this scenario, objective of sustainable yield increase in Bt cotton will drive the major research priorities and policy actions in India.

## Limitations of our study

Partial factor productivity is a reliable estimate for understanding the long-term changes in the yield trends of crops (Fixen et al, 2015, Zhang et al, 2017). However, only three factors namely fertilizer, Bt cotton area, and insecticides were taken in our study for understanding the yield trend in cotton from 2000-14. However, spacing of Bt hybrids, crop management practices, and irrigation measures also showed greater variability at regional levels in the cotton growing regions of India. Thus, comprehensive analysis incorporating agronomic practices coupled with high regional resolution in the data analysis might provide us a highly reliable information on the various factors involved in the observed yield trends of Bt cotton in India.

---

**Supplementary Fig. 1.**
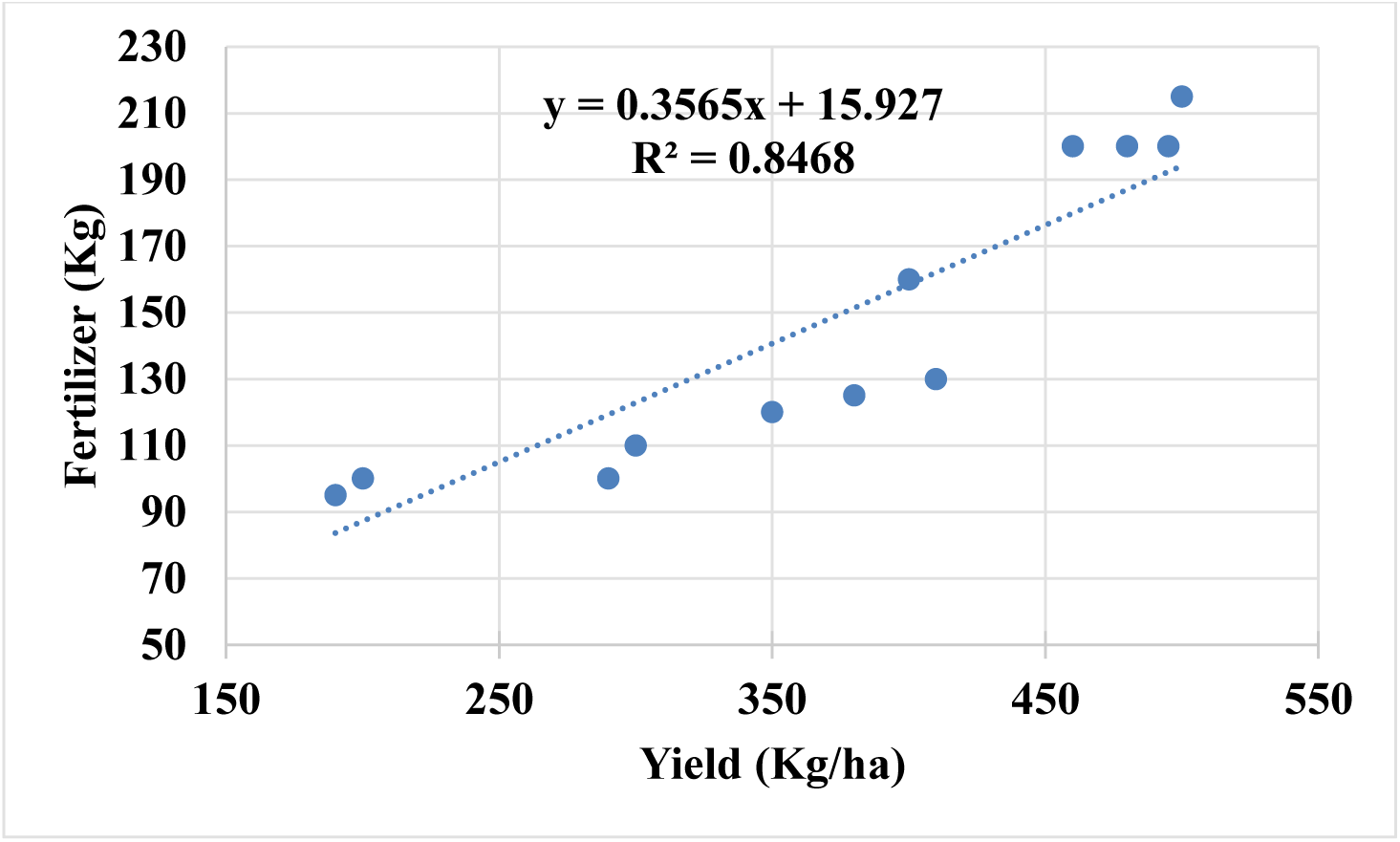
Linear regression of yield and fertilizer usage in cotton during 2000-14

**Supplementary Fig. 2.**
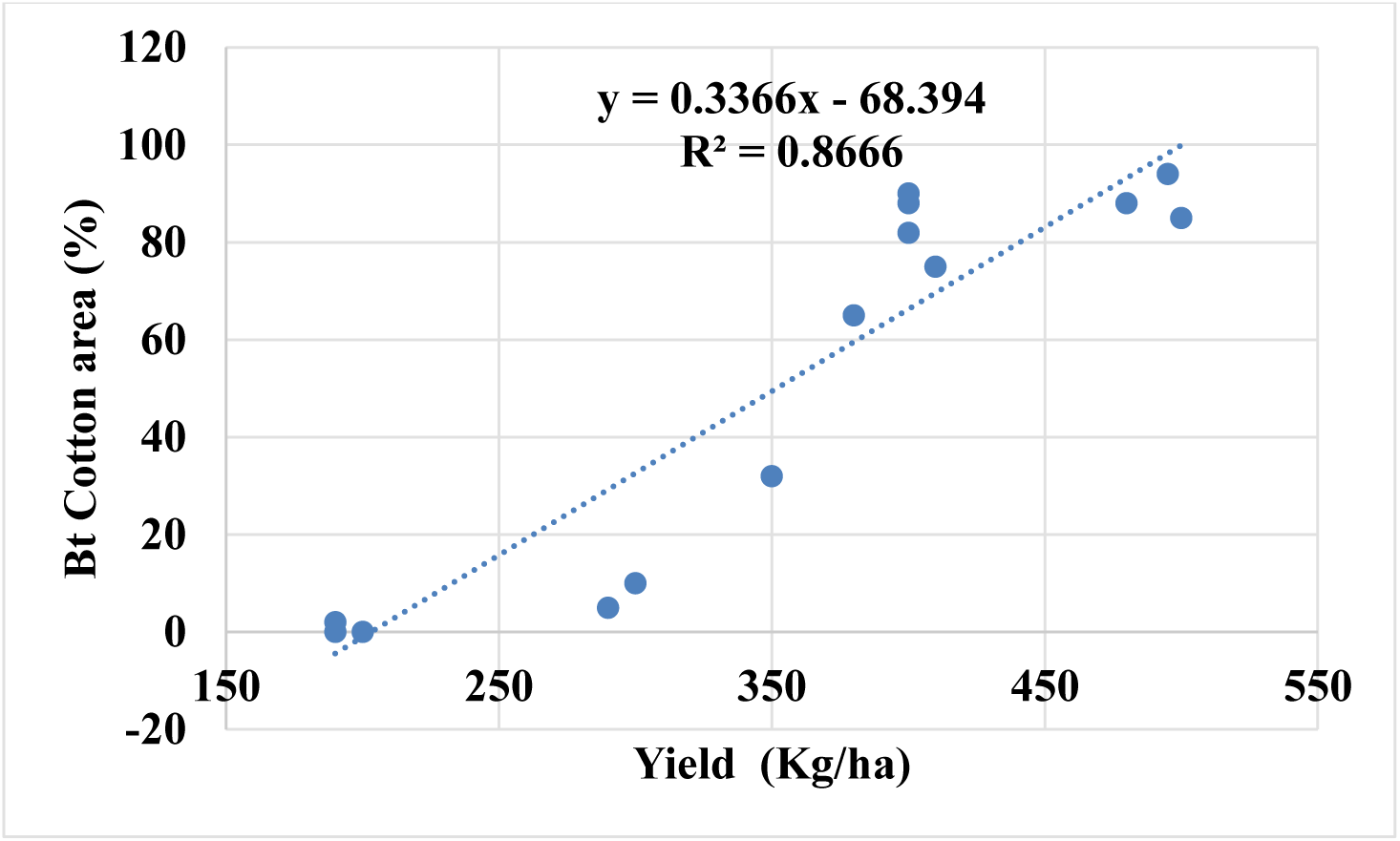
Linear regression of yield and Bt cotton area during 2000-14

**Supplementary Fig. 3.**
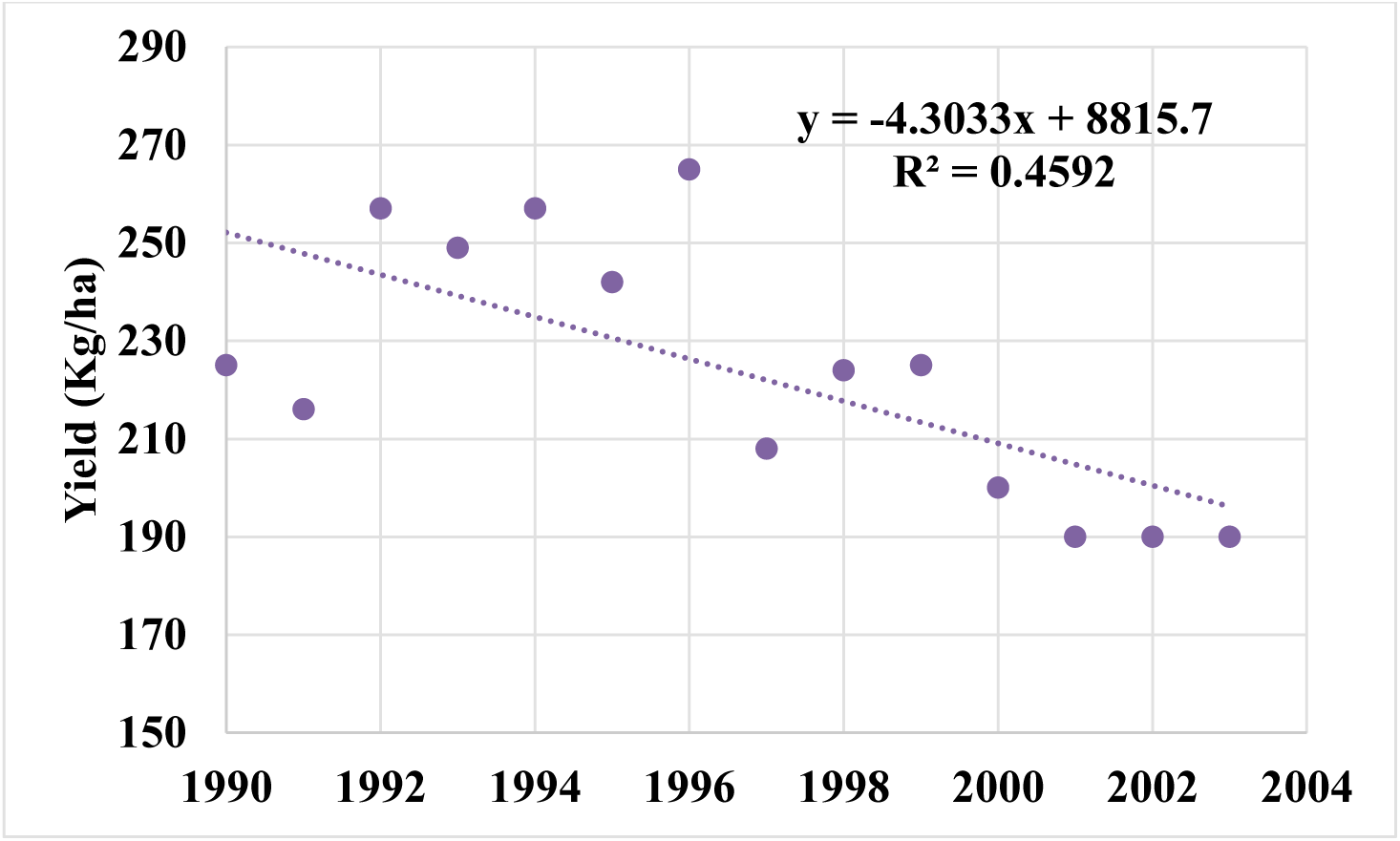
Yield trend of Cotton during 1990-2003

**Supplementary Table 1.** Yield, fertilizer usage and Bt cotton area in India

|  | Yield<br>(Kg/ha) | Bt Cotton<br>area (%) | Fertilizer<br>(Kg/ha) |
| --- | --- | --- | --- |
| 2000 | 200 | 0 | 100 |
| 2001 | 190 | 0 | 95 |
| 2002 | 190 | 0 | 95 |
| 2003 | 190 | 2 | 95 |
| 2004 | 290 | 5 | 100 |
| 2005 | 300 | 10 | 110 |
| 2006 | 350 | 32 | 120 |
| 2007 | 380 | 65 | 125 |
| 2008 | 410 | 75 | 130 |
| 2009 | 400 | 82 | 160 |
| 2010 | 400 | 88 | 160 |
| 2011 | 495 | 94 | 200 |
| 2012 | 480 | 88 | 200 |
| 2013 | 500 | 85 | 215 |
| 2014 | 460 | 90 | 200 |

**Supplementary Table 2.** Yield of Cotton during 1990-2003 in India

| Year | Yield<br>(Kg/ha) |
| --- | --- |
| 1990 | 225 |
| 1991 | 216 |
| 1992 | 257 |
| 1993 | 249 |
| 1994 | 257 |
| 1995 | 242 |
| 1996 | 265 |
| 1997 | 208 |
| 1998 | 224 |
| 1999 | 225 |
| 2000 | 200 |
| 2001 | 190 |
| 2002 | 190 |
| 2003 | 190 |

**Supplementary Table 3.**
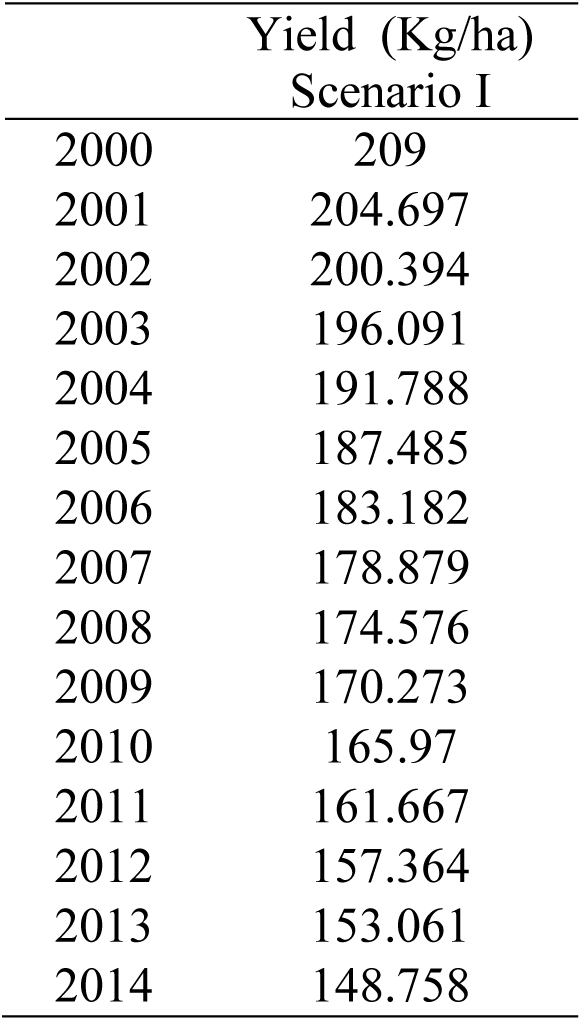
Estimated yield of Cotton in Scenario I

**Supplementary Table 4.** Estimated yield of Cotton in Scenario II

|  | Yield(Kg/ha)<br>Scenario II |
| --- | --- |
| 2000 | 200 |
| 2001 | 190 |
| 2002 | 190 |
| 2003 | 190 |
| 2004 | 200 |
| 2005 | 220 |
| 2006 | 240 |
| 2007 | 250 |
| 2008 | 260 |
| 2009 | 263.68 |
| 2010 | 263.68 |
| 2011 | 329.6 |
| 2012 | 329.6 |
| 2013 | 354.32 |
| 2014 | 329.6 |

**Supplementary Table 5.**
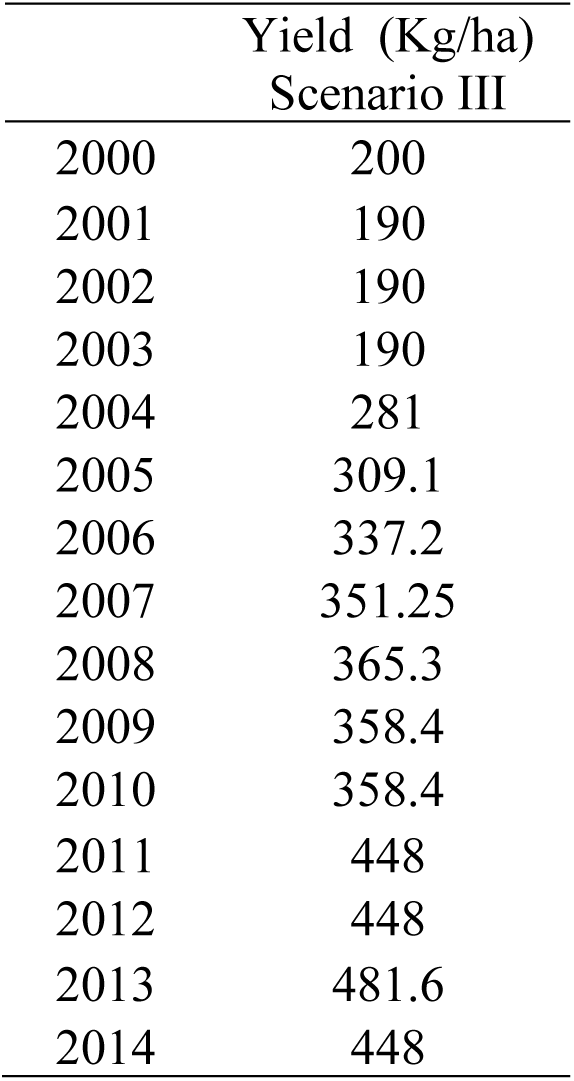
Estimatedyield of Cotton in Scenario III

**Supplementary Table 6.** Quantity (in tonnes) of insecticides used for the control of pests in cotton in India

|  | 2002-03 | 2003-04 | 2004-05 | 2005-06 | 2006-07 | 2007-08 | 2008-09 | 2009-10 | 2010-11 | 2011-12 |
| --- | --- | --- | --- | --- | --- | --- | --- | --- | --- | --- |
| Sucking pest | 2110 | 2909 | 2735 | 2688 | 2374 | 3805 | 3877 | 5816 | 7270 | 6372 |
| Bollworm | 4470 | 6599 | 6454 | 2923 | 1874 | 1201 | 652 | 500 | 249 | 222 |
| other pest | 283 | 537 | 178 | 302 | 375 | 536 | 528 | 410 | 366 | 234 |
| Total | 6863 | 10045 | 9367 | 5914 | 4623 | 5543 | 5057 | 6726 | 7885 | 6828 |

